# Genome-wide Viral Nascent RNA Sequencing Unveils Polymerase Pausing Landscape at Single-nucleotide Precision

**DOI:** 10.1101/2025.01.08.631631

**Authors:** Zhitong Zhu, Cheuk Wang Fung, Xinzhou Xu, Zhibin Liang, Zhuoran Gao, Maggie Haitian Wang, Mingyuan Li, Sin Yu Yeung, Jiguang Wang, Carmen Ka Man Tse, Peter Pak-Hang Cheung

## Abstract

Understanding viral replication and transcription mechanisms is critical for developing effective antiviral strategies. The study of viral gene regulation in host cellular environment is an important bridge for translating mechanistic discoveries from *in vitro* studies to *in vivo* but it remains stagnated due to the absence of technological advancement. Current methods for studying viral transcription and replication have been limited to capturing only the mature viral RNAs, obscuring the dynamic intermediates and mechanistic details of these crucial processes. Here, we introduce an original technology we called “Total Elongating Nascent VIral Polymerase single-molecule Sequencing (TenVIP-seq)”, which isolates and analyses newly synthesized RNA within the viral RNA-dependent RNA polymerase (RdRp) complex, enabling the discovery of mechanisms critical for both viral replication and transcription. Our first characterization of nascent RNA species for an RNA virus showed that RdRP exhibited non-random pausing with profile signatures along the eight influenza A virus (IAV) gene segments. We also revealed genome-wide pausing at known regulatory sites, such as the poly(U) tract polyadenylation site, and new putative regulatory sites at single-nucleotide resolution. Distinct pausing features between the viral genomic (vRNA) and anti-genomic (cRNA/mRNA) templates were observed, suggesting that RdRp processes transcription and replication differentially on positive and negative sense RNA. The NTP analog drug T-705 (favipiravir) intensified RdRp pauses during genome replication and transcription without introducing new pausing sites, while TRIM25 knockout in host cells infected with virus reduced RdRp pausing globally across the viral genome. Strikingly, we observed that terminal nucleotide misincorporations of nascent RNA sequencing of paused RdRP, which were previously undetectable, were as high as an average of 51.6% (vRNA) and 44.0% (cRNA/mRNA), suggesting that the mutational rate of viruses is much higher than previously thought. We demonstrated that TenVIP-seq holds potential in providing insights into the molecular mechanisms that control viral genome replication, gene expression and regulation, mutation, antiviral drug treatment, and virus-host interaction.

## Main

In the five kingdoms of life, cells perform DNA replication and RNA transcription as separate processes^1^. DNA replication guarantees the accurate duplication of the genome, whereas RNA transcription interprets genetic information to synthesize the proteins that facilitate cellular functions^2^. In RNA viruses, both replication and transcription are regulated by a common central apparatus, the RNA-dependent RNA polymerase (RdRp). Despite efforts to identify viral and host factors that mediate replication and transcription^3–5^, how these two fundamental processes are regulated at the genome level in the virus remains elusive. The nature of pausing in viral polymerases, whether stochastic or structured, remains uncertain.

Understanding the molecular mechanisms of viral replication and identifying targets to develop antivirals is critical to cope with new pandemic threats such as Influenza A virus (IAV) and coronaviruses. IAV possesses an eight-segment, single-stranded, negative-sense RNA genome^6^. The viral RNA (vRNA) serves as the template for viral genome replication and transcription. It employs complementary RNA (cRNA) as an intermediary for vRNA replication and conducts transcription via a cap-snatching mechanism. The RdRp, consisting of PB1, PB2, and PA, forms the viral ribonucleoprotein (vRNP) complexed with the viral RNA and NP^7^. The viral mRNA transcription is initiated when the viral RdRp utilizes a host-derived primer obtained through a cap-snatching mechanism^8–10^. Later in the transcription process, the viral RdRp recognizes the poly(U) tract in the viral RNA template and uses it to add the poly(A)-tail to the 3’ end of the viral mRNA, a process known as polyadenylation or transcription termination^11–13^. However, the replication process adopted a *de novo* synthesis approach for both cRNA and vRNA, which requires no priming on the initiation^9,11,12^.

Recent structural and biochemical studies have provided valuable insights into RdRp behaviours in viral transcription and replication, such as stuttering, pausing, and backtracking^14^. For instance, in IAV, the viral polymerase has been shown to stutter *in vitro* during mRNA synthesis at consecutive uracil bases at the 5’ end of the vRNA template^15,16^. Recent data also showed that RdRps in some viruses pause during transcription initiation and backtrack during replication^17^. While non-viral RNA polymerase pausing and backtracking have been demonstrated to play essential roles in transcriptional regulation in eukaryotic and prokaryotic cells^18^, the existence and significance of structured viral RdRp pausing in viral replication, pathogenesis, host cell interactions, and other biological phenomena remains elusive. The interplay between RdRp and the viral RNA molecule during replication and transcription processes, at the cellular or genome-wide level, is yet to be resolved^14^.

RNA-seq enables numerous discoveries about genome organization of viruses, but mechanistic information about gene regulation obtained from these analyses alone is limited as they are unable to provide robust quantification of alternations in RNA synthesis and detection of transient RNA species^19,20^. Nascent RNA sequencing offers solutions to address these limitations. By capturing elongating pre-RNAs associated with RdRp complexes, nascent RNA sequencing can directly observe viral polymerase activity with single-nucleotide precision across entire viral genomes. Also, preserving RNA structures allows for high-sensitivity and high-resolution detection of splicing programs and regulatory elements. Meanwhile, studying the effects of host factors on polymerase dynamics across the viral genome provides novel insights into viral-host interplay during viral replication and transcription. Lastly, distinguishing nascent RNA from processed and mature transcripts provides precise temporal profiling of changing expression patterns. Observing initiation, pausing and termination sites reveals previously unknown mechanisms of transcriptional control. Therefore, nascent viral RNA sequencing, which captures the dynamic process of viral transcription and replication within infected cells, serves as a powerful tool to translate *in vitro* structural biology and enzymology observations into a realistic understanding of the complex mechanisms underlying viral infections in the real physiological cellular environment.

To date, applications of nascent RNA sequencing to study virus-related topics have exclusively restricted on the host’s transcriptomic changes following viral infection^21–23^ and do not investigate viral replication and transcription through viral polymerases. Also, viral replication and transcription do not follow eukaryotic counterparts, such as template switching^24^, rolling-circle replication^25^, uncoupled replication and transcription^26^, and exploitation of host factors^27^. Hence, nascent RNA sequencing can unravel many hidden but transient replication and transcription processes that current mature RNA sequencing techniques failed to capture. Overall, the development of a nascent RNA sequencing platform customed engineered for virus offers the potential in uncovering many viral gene regulation processes by elucidate the dynamics of viral replication and transcription represents an unmet need constituting a major setback in fundamental biology.

NTP analog is one of the most important classes of antiviral drugs in defensing overwhelming pandemics. Plenty of work has been done in describing the mechanism of actions of these analogs on the viral polymerase *in vitro*, as well as the analog efficacy on virus in cell experiments. However, translating outcomes from different levels remain challenging for the NTP analog drugs, because the mechanism of action of NTP analogs drugs on RdRp cannot be directly recapitulated in the genome-wide cellular level^14^. It is unfeasible for structural biology, enzymology, and virology studies to bridge the gap between single analog molecule incorporation event and the effects on pan-genomic context.

How host factors impact RdRp replication and transcription genome-wide inside cells have not been directly studied due to the absence of tools. For example, tripartite motif-containing protein 25 (TRIM25) is a host protein that binds to the influenza vRNP in the nucleus of infected cells. TRIM25 acts as a molecular clamp on the vRNPs, preventing the viral polymerase from accessing the viral RNA templates^28^. This inhibits viral RNA chain elongation, without affecting the initiation of viral mRNA synthesis. The exact mechanism by which TRIM25 hinders the IAV polymerase from elongating the RNA chain is not fully elucidated. This is due to the lack of tools and techniques that can directly probe the dynamics of the viral RdRp complex and its interactions with host factors in the native cellular environment.

The IAV RdRp is an error-prone enzyme with no proofreading mechanism. Often, it leads to high mutation occurrences within the virus genome^29^. This mechanism plays a major role in escaping the population immunity due to the rapid change in the viral surface glycoproteins hemagglutinin and neuraminidase, a process known as antigenic drift. Polymerase infidelity directly modulates viral mutation rates^60^. Misincorporations are also contributed by other factors, including sequence context, cellular microenvironment, replication mechanisms, proofreading, and access to post-replicative repair^3,30,31^. However, how the cellular genome-wide dynamic activity of viral polymerase changes once misincorporation are made has not been studied. Elucidating the polymerase dynamics while copying the viral genome during misincorporation can help to explain the fundamental mechanisms of RNA synthesis and mutation, highlight the critical structural features and kinetic checkpoints that enable polymerases to discriminate between correct and incorrect substrates, and explain the origins of these errors and their biological and evolutionary consequences.

Investigating viral RNA synthesis mechanisms via nascent RNA sequencing would necessitate multifaceted innovative methodologies custom-tailored to the unique molecular biology of viral polymerases. Current IAV RNA sequencing methods focusing on the whole genome of packaged and released IAV particles are unable to capture active transcription and replication events, which occur inside the nucleus. Termination and end labeling by the incorporation of biotin-labeled nucleotide analog as described in Pro-Seq failed to halt or get incorporated into the viral RNA synthesis efficiently in viral RdRp. Unlike the eukaryotic nascent RNA sequencing technology, currently, there is no approach to capture and examine the transient nascent RNA species of RNA viruses during active transcription or replication events. The difficulties may come from the paucity of methods to effectively enrich the active viral RdRp and detect the elongating viral RNA on the RdRp. To overcome all these challenges, we present Total Elongating and Nascent VIral RNA Polymerase single-molecule RNA sequencing (TenVIP-seq), an IAV nascent RNA-sequencing method, featuring a PCR-free approach that enriches and profiles all viral nascent RNA species from infected cells via DRS. In the context of eukaryotic system, nascent RNA only refers to the mRNA transcript, but for RNA virus system, the nascent RNA refers to the viral mRNA, vRNA, and cRNA replicons and transcripts. Hence, we designed a novel library preparation method for Nanopore sequencing to capture all viral nascent RNA that are still bound on the IAV RdRp. This method enabled us to identify various pausing at gene regulatory sites during viral RNA replication and transcription. Such paused polymerases provide snapshots of the replication regulation in action. TenVIP-seq uncovered many new biological insights, centering on gene regulation, pathogenesis, antiviral treatment, and virus-host interactions. Our techniques warrant much future work to further examine in detail the new biological insights we reported here.

### Isolation and Sequencing of Influenza A Virus Nascent RNA

To capture the full IAV heterotrimeric RdRp-RNA complex that is actively transcribing RNA, we first generated recombinant virus carrying affinity tag for purification. A streptavidin tag (strep-tag) was inserted into the A/WSN/1933(H1N1) (WSN) PB2 vRNA sequence of the full RdRp to achieve pull-down of the RdRp-RNA complex (**Fig. S1a**). Plasmids with the PB2 with the strep-tag modification at C-terminal, and the remaining seven plasmids with the WSN gene segments PB1, PA, HA, NP, NA, M, and NS were used to generate the complete viral particles with PB2 carrying a strep-tag (WSN PB2-strep). This modification allowed the specific enrichment of the RdRp complexes, together with the actively synthesizing RNA and its template from the nuclei of host cells infected by the WSN PB2-strep virus. To determine if the addition of strep tag to the viral genome affects viral replication, plaque assays were used to show that both WSN wild-type and WSN-PB2 strep replicated to similar plaque size (**Fig. S1b)**. RT-qPCR of purified RNA from infected cells showed similar NP gene expression between two viruses (**Fig. S1c**). Lastly, the virus growth kinetics from 3 to 36 hours post-infection (h.p.i.) indicated comparable viral replication efficiencies between the wild-type strain and the PB2-strep WSN variant (**Fig. S1d**). Based on reported studies^32–34^, 4-hour post-infection at the multiplicity of infection (M.O.I.) of 1 was employed.

To evaluate TenVIP-seq, human alveolar basal epithelial cells (A549) were infected with the WSN PB2-strep virus. As IAV replicates and transcribes within the host nucleus, the cell nuclei were extracted to perform TenVIP-seq (**Fig. 1a**). During library preparation, multiple checkpoints were employed to assess the integrity of the samples. First, the presence of PB2 and the strep-tag were confirmed in both nuclei and cytoplasmic lysates, with histone H3 and tubulin serving as markers for lysate origins, respectively (**Fig. 1b**). Further western blot analysis confirmed the presence of PB2, strep-tag, PB1, and NP in the nuclei lysate input and elute. Notably, the biotin column selectively enriched for RdRp complexes containing the PB2-strep subunit, showing no RdRp complexes in the elute from WSN wild-type (**Fig. S1e**).

**Fig 1:**
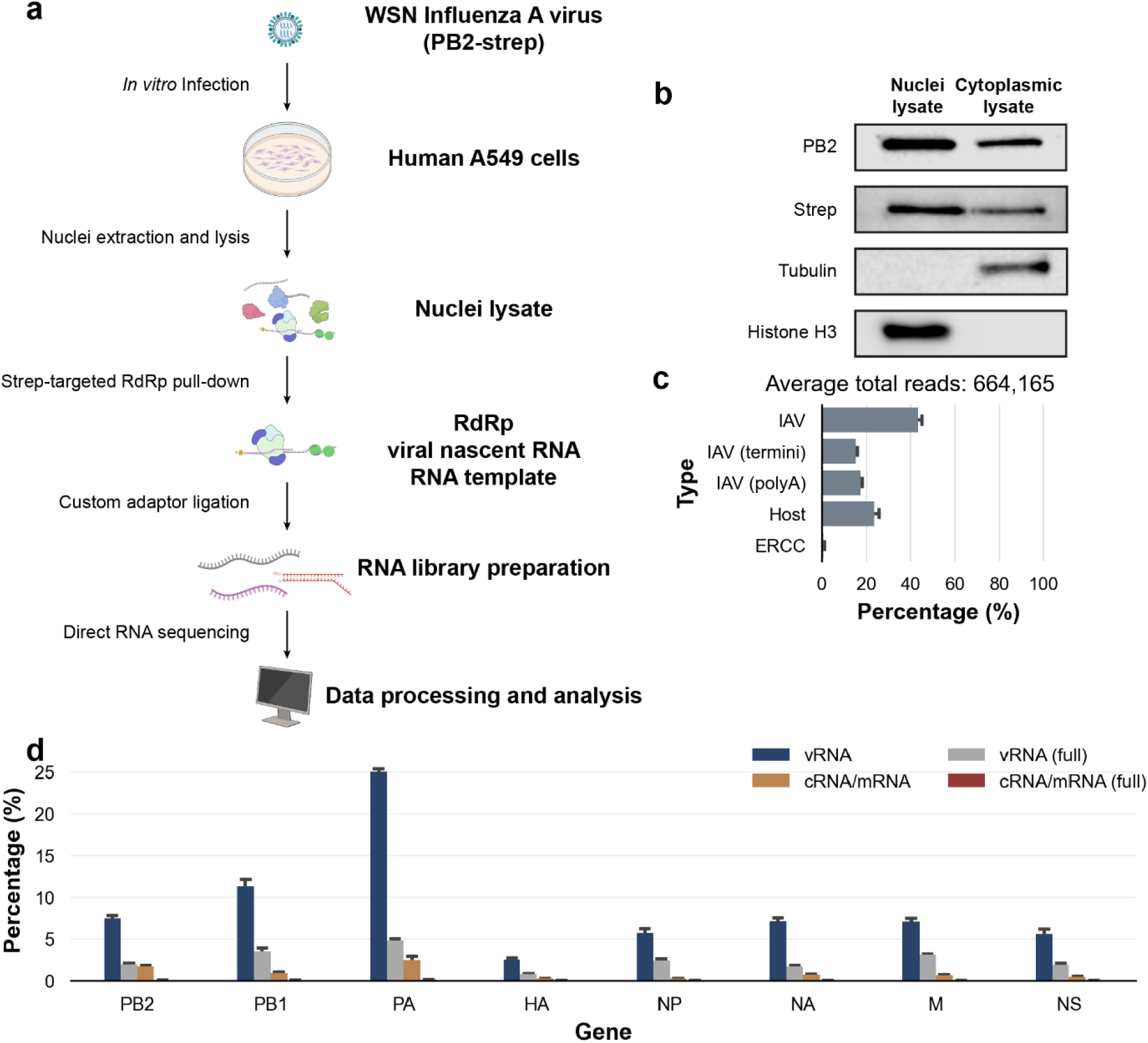
Capture and RNA Sequencing of Influenza A Virus Using TenVIP-seq. **a.** Schematic representation of TenVIP-seq with PB2-recombinant WSN, utilizing DRS with a modified DNA adaptor. **b.** Western blot analysis of lysates during TenVIP-seq library preparation, targeting PB2, Strep-tag, tubulin, and histone H3. **c.** Percentage of each RNA species in uniquely mapped reads, presented as mean ± SEM, n = 3. **d.** Percentage of vRNA and cRNA/mRNA reads for each IAV gene, including reads aligned to the 3’ end of the respective RNA species. Data presented as mean ± SEM, n = 3.

Unlike poly(A)-tailed mature mRNA, nascent RNA lacks a generic sequence that could be utilized for enrichment and sequencing library preparation. Given that Nanopore DRS is designed for mature mRNA, a novel strategy for incorporating sequencing adaptors to the viral nascent RNA is essential. To address this, we developed a customed-engineered ligation approach featuring a new adaptor design. To mitigate residual polymerase activity during pull-down, Zn^2+^ was added when the nucleus was isolated on ice, effectively inhibiting IAV RdRp activity (**Fig. S2**). This step ensures that the polymerase positions identified in the data analysis accurately reflect their locations during replication and transcription within the host nucleus.

Our sequencing data demonstrated successful enrichment and sequencing of nascent RNA from the WSN PB2-strep (**Fig. 1c**, n = 3), with over 40% of reads aligning to the IAV genome. Notably, only 17.4% of these reads contained a poly(A)-tail, suggesting the capture of mature mRNA. While the presence of RNA templates for synthesis and RNA in the newly packaged vRNP and cRNP complexes were expected in TenVIP-seq due to the association of the RdRp and the template/genome^9^, 15.1% of reads aligned to the last base of the reference, indicating the potential presence of full-length RNA template during RNA synthesis and strep-tag pull-down. 23.4% of reads aligned with the human transcriptome, reflecting host-derived RNA. The distribution of reads between vRNA and cRNA/mRNA strands, which share the same directionality, favored the production of vRNA species across all eight segmented genomes (**Fig. 1d**). RNAs corresponding to RdRp subunits, specially PB2, PB1, and PA, were among the most abundant at 7.4%, 11.3%, and 25.1% respectively, while HA, NP, NA, M, and NS contributed 2.5%, 5.7%, 7.2%, 7.1%, and 5.6% respectively (**Fig. 1d**). Additionally, read coverage of ERCC spike-in mix also confirmed that TenVIP-seq effectively captures RNA molecules of varying sizes with a strong correlation (**Fig. S3a**, r = 0.99), demonstrating a robust dynamic range.

### Direct Evidence of RdRp Pausing Signatures and Pausing at Regulatory Sites

Utilizing TenVIP-seq, we accurately mapped the positions of the RdRp complex during the syntheses of vRNA, cRNA, and mRNA, providing a genome-wide view of both genome replication and transcription of the WSN at single nucleotide resolution (**Fig. 2a, c**). Nascent RNA reads were detected across the vRNA and cRNA genomes, with reads also consistently present at the 3’ terminal of both vRNA or cRNA/mRNA. We applied the same bioinformatic pipeline we developed for TenVIP-seq to the A/Puerto Rico/8/1934 (H1N1) (PR8) dataset^35^ to validate our hypothesis that the mapped 3’ termini of the nascent viral RNA indicated genuine polymerase pausing. These results suggested that the observed profiles originated not only from mature RNA templates but also from nascent RNA. Similar coverage at the 3’ end of the vRNA was observed (**Fig. 2b**), suggesting that the read coverages in our dataset indeed represented RNA templates. Despite the presence of full-length templates, nascent RNA coverage was evident within the genome segments, and the distinct 3’-biased distribution of mature vRNA facilitated differentiation of pausing profiles identified by TenVIP-seq from template and mature RNA.

**Fig. 2:**
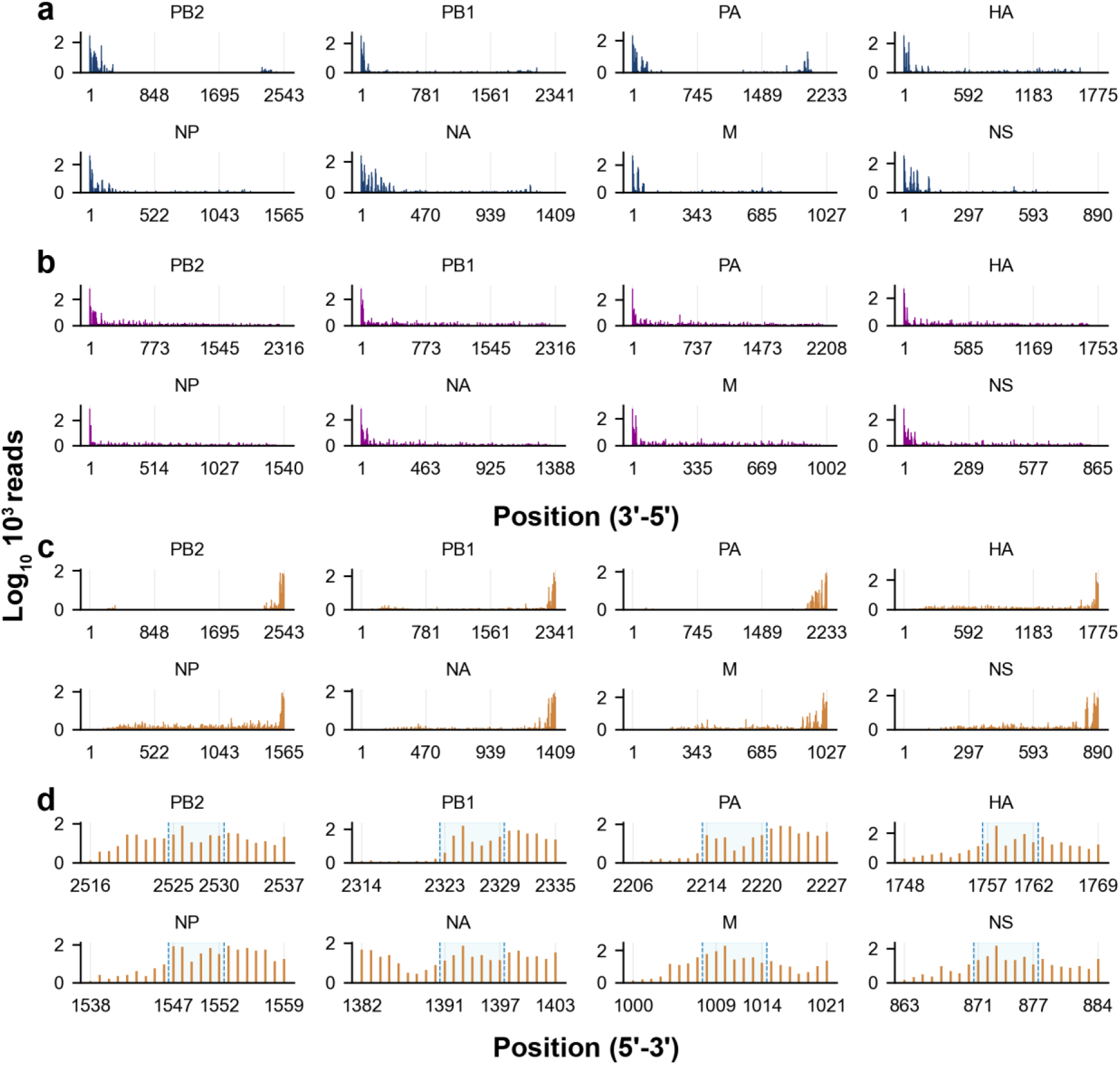
The last nucleotide coverage of RNA molecules from WSN PB2-strep (n = 3) across the eight-segmented genome is compared to PR8 as a reference for mature vRNA. **a.** Last nucleotide coverage of WSN nascent vRNA across the complete genome. **b.** Last nucleotide coverage of mature vRNA from PR8 (from public dataset^35^) **c.** Last nucleotide coverage of WSN nascent cRNA/mRNA across the genome. **d.** Position of WSN RdRp at the U track of the template, marking the initiation of poly(A) tail synthesis. U track positions for each gene are highlighted.

When the RdRp encounters the poly(U) tract on the vRNA template, enzymology studies showed that it stutters at this site, efficiently incorporating the corresponding adenine residues to synthesize the poly(A) tail of mature viral mRNA^15^. However, it remains unclear whether these observations accurately reflect polymerase dynamics within the physiological intracellular environment and how they influence RdRp activity. Examination of the 3’ regions of cRNA and mRNA revealed significant pausing signals within a 5-6 nucleotide stretch of the poly(U) track (**Fig. 2d**). This finding suggests that the TenVIP-seq approach effectively captures RdRp pausing events during mRNA polyadenylation by the viral RdRp, enabling further investigation into this known stuttering mechanism during viral RNA transcription. The ability to capture sequence-specific regulatory sites demonstrated that TenVIP-seq is both sensitive and precise at single-nucleotide resolution.

TenVIP-seq demonstrates high reproducibility between technical and biological replicates, suggesting that RdRp pausing in viruses is not completely stochastic. Using the WSN PB2-strep virus, the pausing profiles of both vRNA (**Fig. 3a**, top panels) and cRNA/mRNA (**Fig. 3b**, top panels) from technical repeats showed no observable differences, while only minor significant differences were noted between biological repeats (**Fig. 3a, b**, bottom panels, **Fig. S4**). These findings indicate that the TenVIP-seq approach consistently generates and highly reproducible pausing profiles of the viral RdRp across technical replicates. Importantly, the robustness of the observed pausing signatures suggests that the pausing event of viral polymerase is not stochastic, but instead reflects genuine regulatory dynamics. This reinforces the reliability of this method in capturing the true activity of RdRp during viral genome replication and transcription. In summary, this accurate, high-resolution approach enables quantitative assessment of RdRp density at viral genome segments, serving as a proxy for potential pausing events that indicate viral or host regulatory processes such as poly(U) track. Moreover, TenVIP-seq can be leveraged to study alterations in the pausing landscape under various experimental conditions, such as viral gene mutations and host adaptations, using statistical analysis to comprehensively identify and compare genomic sites that regulate RdRp dynamics.

**Fig. 3:**
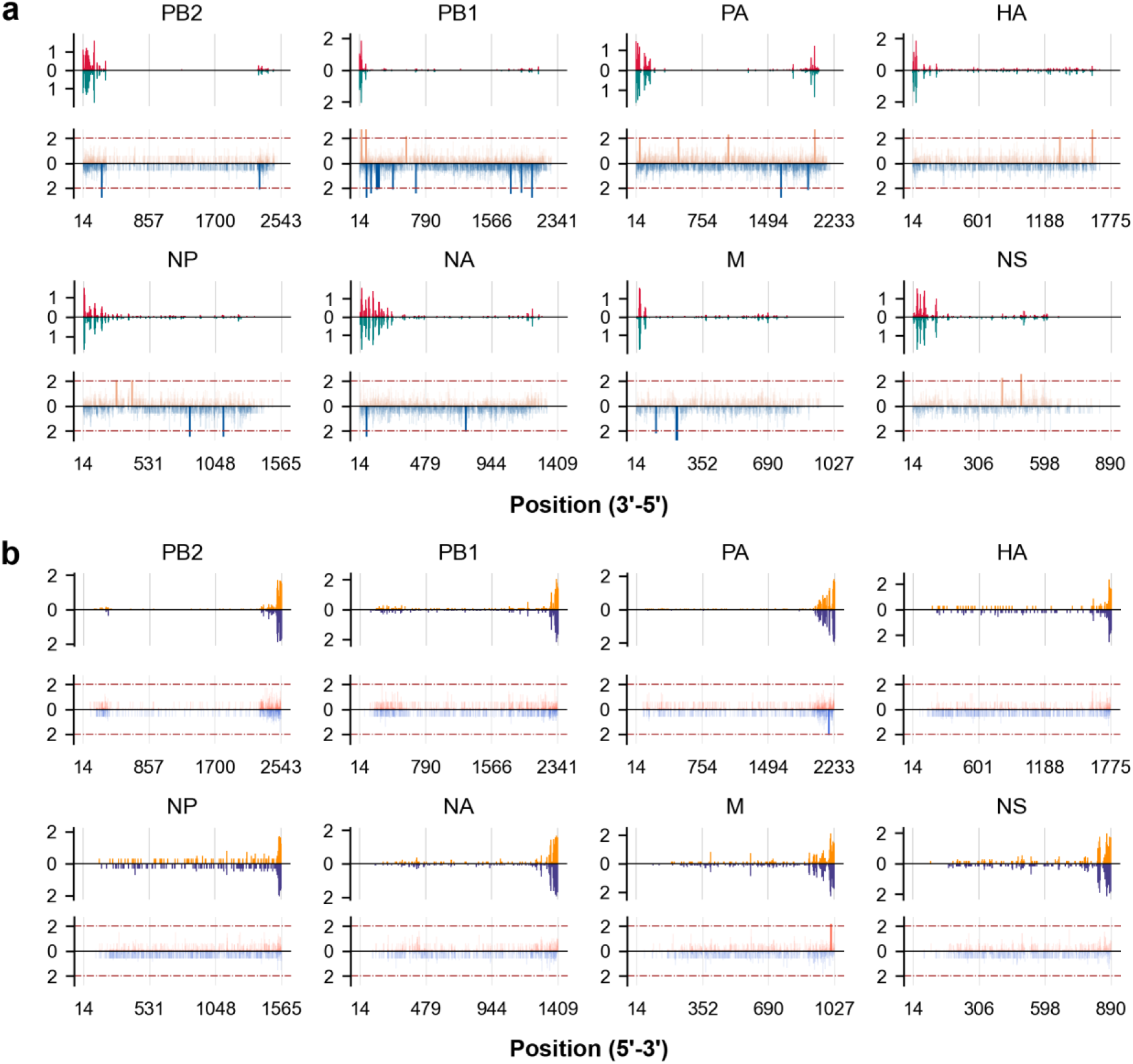
Genome-wide view of WSN PB2-strep last nucleotide coverage. Top: Comparison of last nucleotide coverage between technical replicates (red and green). Bottom: Statistical significance (-log_10_(*p*-value)) between biological replicates (orange and blue, n = 2). Statistical differences in coverage at each position were calculated using Welch’s t-test. **a.** Coverage view and statistical differences for WSN vRNA segments. Dotted lines at -log_10_(*p*-value) = 2 indicate a *p*-value of 0.01. **b.** Coverage view and statistical differences for WSN cRNA/mRNA segments. Dotted lines at -log_10_(*p*-value) = 2 indicate a *p*-value of 0.01.

### Genomic effects of viral RdRp replication and transcription by NTP analog drugs

Assessing the effectiveness of NTP analog drugs presents significant challenges, particularly in extrapolating *in vitro* enzymology data to a comprehensive cellular context on a genome-wide scale. The effects of these NTP analog drugs on viral polymerase function, as well as their impact on viral replication and transcription across the entire viral genome, remained underexplored due to a lack of robust methodologies. Previous research in this field has been limited, primarily relying on *in vitro* polymerase assays, cell-based antiviral studies, and resistance mutation analysis^36,37^. While these approaches are informative, they do not evaluate the drug’s effects on its target enzyme within the context of the full viral genome under physiological conditions. Genomic studies of NTP analogs have largely focused on their role in generating drug resistance in the virus, without direct examination of the direct impact on the target enzyme in a realistic physiological setting on a genome-wide scale. The TenVIP-seq approach offers a solution to bridge this gap in enzymology and genomics. It enables an in-depth, systems-level investigation into the mechanisms of action of NTP analog drugs, elucidating how they disrupt the coordinated functioning of the viral polymerase complex and host proteins across the entire viral genome.

We first investigated the impact of the NTP analog drug T-705 (favipiravir) on the relative levels of newly synthesized RNA expression across the eight IAV gene segments. T-705 was administrated to the culture medium of A549 cells for 9 hours, with infection occurring 4 hours prior to the preparation of the TenVIP-seq library (**Fig. 4a**). To confirm that the drug’s effectiveness in disrupting nascent RNA synthesis, we conducted RT-qPCR analysis on both the viral NP gene vRNA and cRNA from RNA purified from infected cells treated with 10 and 1000 µM T-705. Notably, the levels of both vRNA and cRNA decreased significantly with increasing concentrations of T-705 throughout the experiment (**Fig. 4b**). To gain deeper mechanistic insights, we analyzed the genome-wide pausing profiles of the RdRp complex in the presence of T-705 treatment (**Fig. 4c-h; Fig. S5a-d**). Strikingly, our statistical analysis revealed that T-705 predominantly increased the intensities of existing polymerase pausing sites across the viral genome, as observable at the coverage view (**Fig. 4c-h; Fig. S5a-d**), rather than inducing entirely new pausing locations, indicated by the number of positions with significant difference in normalized pausing counts between T-705 treated cells and untreated control (**Fig. 4d, g** (bottom panels), **Fig. S5e**). These results indicate that the primary mode of action for T-705 is to disrupt the progression of the viral polymerase complex by modulating the frequency and duration of pausing events, rather than causing wholesale changes to the landscape. The TenVIP-seq platform has enabled us to uncover these nuanced, yet crucial, mechanistic details that could inform the rational design of more effective NTP analog therapeutics targeting the viral replication machinery.

**Fig. 4:**
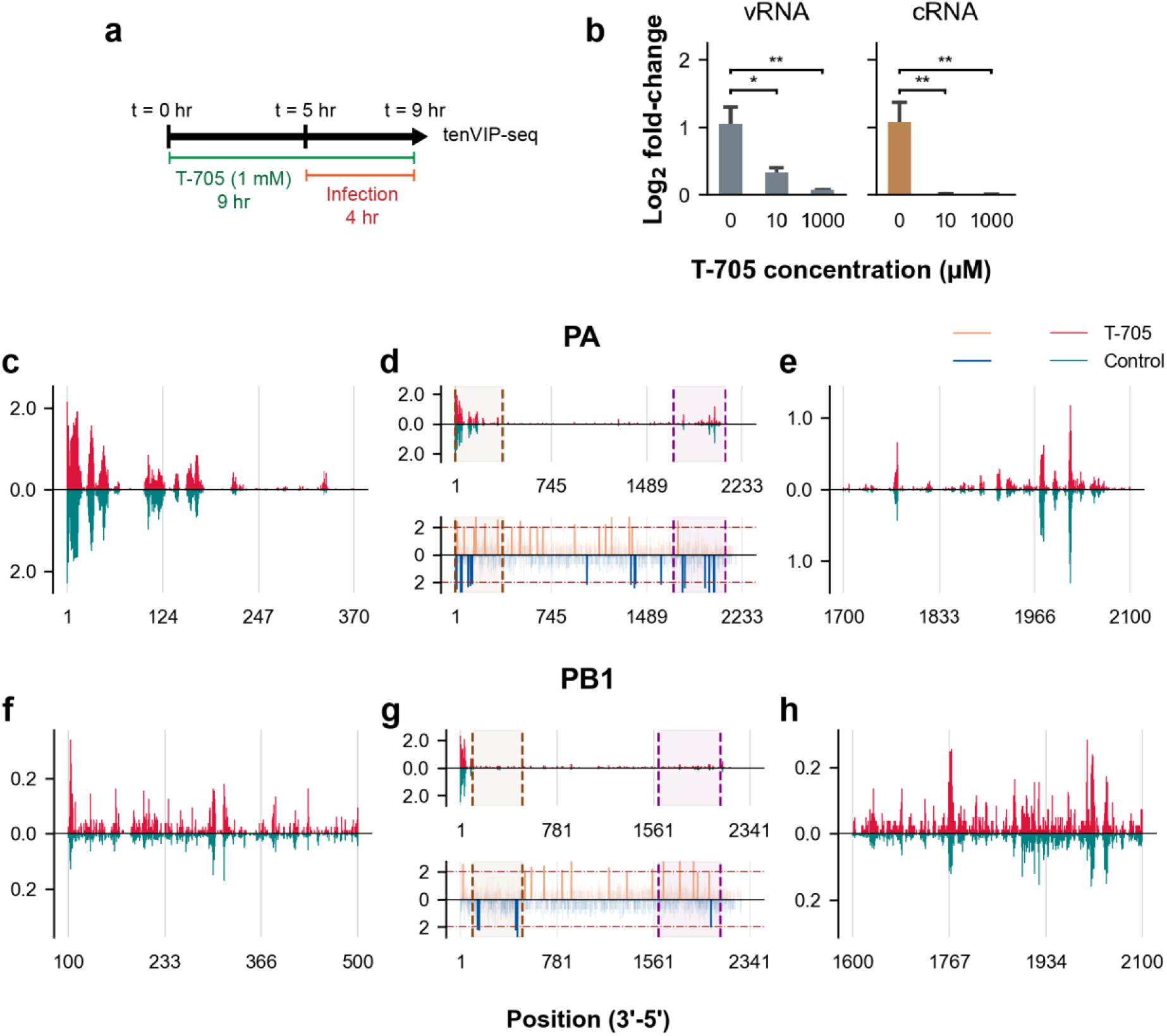
T-705 Increases RdRp Pausing Frequency. **a.** Time course of T-705 treatment (1 mM) on the A549 cell line, followed by TenVIP-seq. **b.** T-705 treatment significantly decreased both WSN vRNA and cRNA expression (One-way ANOVA followed by Dunnett’s multiple comparison test; ANOVA *p*-value < 0.01. **: *p*-value < 0.01; *: *p*-value < 0.05; n = 3). **c-e.** Last nucleotide coverage and statistical significance (Welch’s t-test) of viral PA vRNA between T-705-treated cells (top panel, n = 2) and untreated controls (bottom panel, n = 3), observed in the 3’-5’ direction. **c**. Expanded coverage view of PA vRNA from 1 to 370 nucleotides. **d.** Statistical differences in the normalized pausing count at each genomic position between the T-705 treatment and no treatment groups were calculated using Welch’s t-test. Coverage of full-length PA vRNA (top) and -log_10_(*p*-value) for each nucleotide coverage of PA vRNA (bottom) between the two samples. Dashed lines at -log_10_(*p*-value) = 2 indicate a *p*-value of 0.01. Dotted lines and shading represent the expanded coverage view. **e.** Expanded coverage view of PA vRNA from 1700 to 2100 nucleotides. **f-h.** Last nucleotide coverage and statistical significance of viral PB1 vRNA. **f.** Expanded coverage view of PB1 vRNA from 100 to 500 nucleotides. **g.** Coverage of full-length PB1 vRNA (top) and -log_10_(*p*-value) for each nucleotide coverage of PB1 vRNA (bottom) between the two samples. **h.** Expanded coverage view of PB1 vRNA from 1600 to 2100 nucleotides.

### Host factors alter viral RdRp replication and transcription dynamics

TenVIP-seq may serve as a valuable tool for investigating how host factors regulate viral RdRp transcription and replication across the entire viral genome. Current techniques have been limited to assessing the impacts of host factors on RdRp activity *in vitro* or through endpoint virus titer assays using knockout cell lines or animal models^38^. These approaches provide limited insights into how host factors affect RdRp in the host environment, as *in vitro* data often do not translate effectively to the *in vivo* environment. TenVIP-seq enables direct genome-wide measurement of nascent viral RNA synthesis, capturing RdRp transcription kinetics along each gene segment in infected cells. This systems-level view can elucidate how cellular proteins or other host factors influence RdRp processivity, pausing, or termination events during influenza replication and transcription.

For example, TRIM25 is an E3 ubiquitin ligase that mediates K63-linked ubiquitination of the RIG-I receptor, a critical component for innate immune sensing of influenza virus and induction of type I interferons^39^. Previous studies have shown that TRIM25 restricts influenza virus replication^40,41^. However, the underlying mechanisms remain incompletely understood. TenVIP-seq can determine whether TRIM25 affects influenza RdRp kinetics, such as processivity or pausing, across the viral genome. This would provide insights into how TRIM25 restricts viral replication. Currently, there is an absence of methods to evaluate how host factors like TRIM25 modulate influenza RdRp transcriptional activity directly at the genomic level within infected cells, a limitation that TenVIP-seq overcomes.

In our study, we infected A549 wild-type and TRIM25-knockout (KO) cells with the recombinant IAV at the same M.O.I. of 1 for 4 hours. RT-qPCR analysis revealed elevated expression level of interferon and RIG-I genes, indicating an active innate antiviral response from the host (**Fig. 5a**). Western blot analysis further confirmed that, despite the upregulation of antiviral response genes, TRIM25 protein was not expressed in TRIM25-KO cell line (**Fig. 5b**), validating the effectiveness of the KO even in the presence of increased antiviral gene expression. After system validation, we conducted a detailed comparative analysis of the RNA polymerase pausing profiles across the IAV genome under the two cellular conditions. Specifically, we compared the pausing dynamics of the viral RdRp complex during IAV infection of wild-type cells versus cells with the host factor TRIM25 genetically knocked out (**Fig. 5c-h; Fig. S6a-d**). Strikingly, our analysis revealed that the overall intensity and distribution of RdRp pausing events were significantly reduced across the viral genome during both viral replication and transcription in the TRIM25-KO cells (**Fig. 5c-h; Fig. S6a-d**). The number of positions with significant difference in normalized pausing counts between wild-type and TRIM25-KO cells (**Fig. 5d, g (bottom panels), Fig. S6e**) This finding demonstrates the critical role of the host TRIM25 protein in reshaping the pausing landscape of the viral polymerase complex as it traverses the IAV genome.

**Fig. 5:**
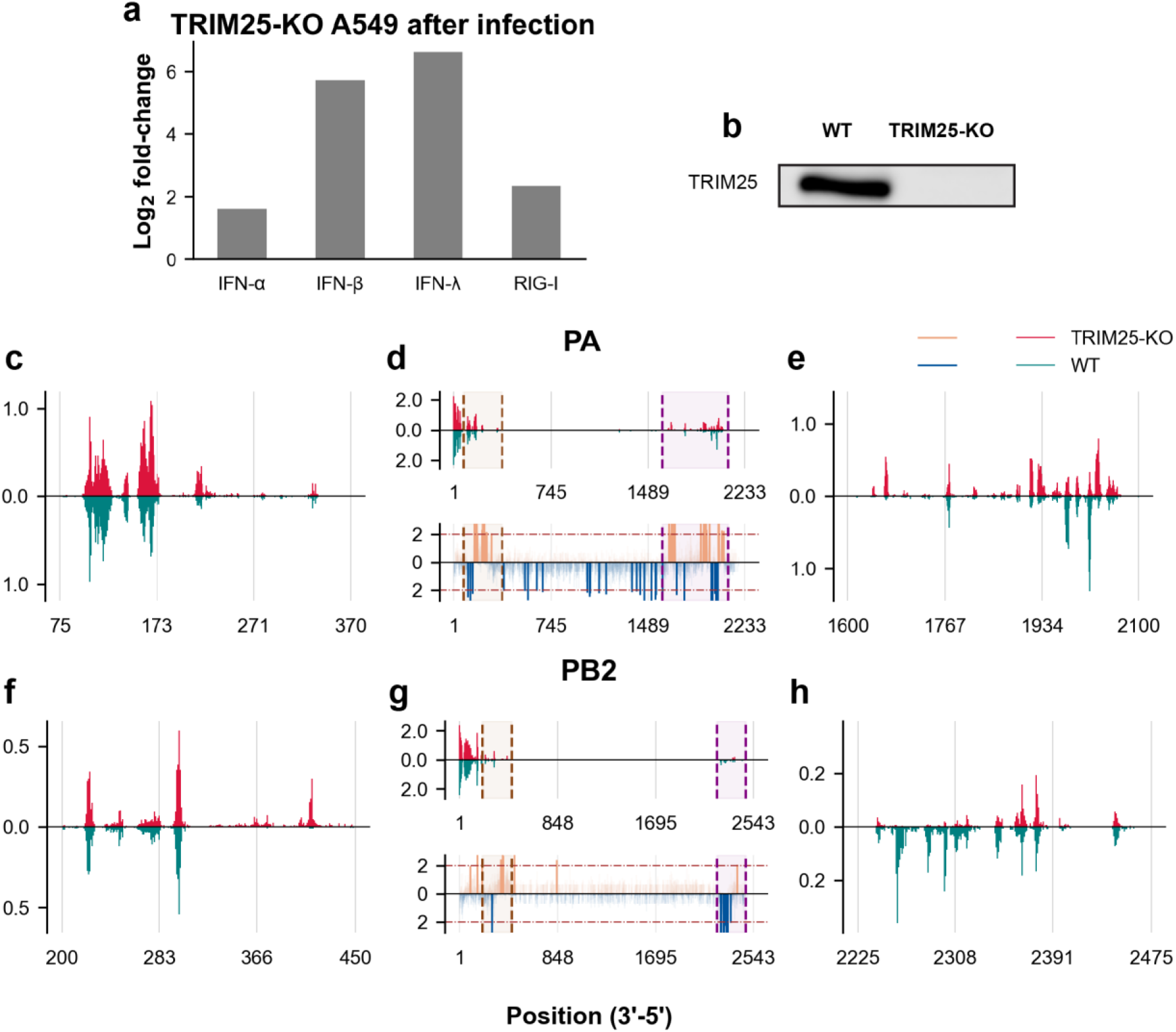
TRIM25 KO Alters Pausing Profile. **a**. Changes in the expression levels of antiviral response genes in the TRIM25 knockout (KO) A549 cell line at 4 hours post-infection (h.p.i.), with fold-change normalized to GAPDH. **b**. Western blot analysis of TRIM25 protein expression in wild-type and KO cell lines. **c-e**. Last nucleotide coverage and statistical significance (Welch’s t-test) of viral PA vRNA between TRIM25-KO cells (top panel, n = 3) and wild-type controls (bottom panel, n = 3), observed in the 3’-5’ direction. **c**. Expanded coverage view of PA vRNA from 75 to 370 nucleotides. **d**. Coverage of full-length PA vRNA (top) and -log_10_(*p*-value) for each nucleotide coverage of PA vRNA (bottom) between the two samples. Dashed lines at -log_10_(*p*-value) = 2 indicate a *p*-value of 0.01. Dotted lines and shading represent the expanded coverage view. **e**. Expanded coverage view of PA vRNA from 1600 to 2100 nucleotides. **f-h**. Last nucleotide coverage and statistical significance of viral PB2 vRNA. **f**. Expanded coverage view of PB2 vRNA from 200 to 450 nucleotides. **g**. Coverage of full-length PB2 vRNA (top) and -log_10_(*p*-value) for each nucleotide coverage of PB2 vRNA (bottom) between the two samples. **h**. Expanded coverage view of PB2 vRNA from 2225 to 2475 nucleotides.

### Pausing is predominantly correlated with nucleotide misincorporation by viral polymerases

When misincorporation events occur during RNA synthesis, RNA polymerases in cell, bacteria, and coronavirus that are capable of proofreading engage in a backtracking process that leads to transcriptional pausing^18^. Recent studies have demonstrated that influenza A and B polymerases stall, backtrack, and chain terminate upon consecutive incorporation of nucleoside analogs^42^. However, the dynamics of polymerases lacking proofreading abilities, as of nearly all viral polymerases except those of coronaviruses, during nucleotide misincorporation in replication or transcription remain poorly understood. To investigate the whole-genome level viral polymerase dynamics upon misincorporation, we analyzed the 3’ end mismatches of nascent vRNA and cRNA/mRNA of WSN PB2-strep using TenVIP-seq. Strikingly, we observed that an average of 51.6% of the vRNA and 44.0% c/mRNA transcripts contained misincorporations at the 3’ terminal of the read, while the remaining 48.4% of vRNA reads and 56.1% of cRNA/mRNA reads with RdRp paused did not exhibit misincorporation (**Fig. 6a**). When considering two consecutive nucleotide misincorporations, AU mismatch at the 3’ end predominantly contributes 70.5% which were incorporated opposite the templating vRNA nucleotide prior to RdRp pausing or termination (**Fig. 6b**, left). Interestingly, while AU misincorporation remained the highest at 19.7%, the types of misincorporations during cRNA/mRNA synthesis were more diverse compared to those observed during vRNA synthesis (**Fig. 6b**, right). In addition, 3 and 4-nt long misincorporation motifs in vRNA synthesis showed AUC and AUCC were dominantly highest in their categories, at the mean percentage of 37.1% and 25.0% respectively (**Fig. S7**, left). Meanwhile, the 3 and 4-nt long mismatch at the cRNA/mRNA 3’end remained diverse compared to vRNA counterpart (**Fig. S7**, right). Further exploration of misincorporation motifs at the 3’ end between vRNA and cRNA/mRNA during synthesis leads to the identification of the most prominent motifs. In vRNA, AU contributed the highest mean percentage of 22.8% of motifs 1-4 nt long (**Fig. 6c**, left); in cRNA/mRNA, AU contributed only 5.6%, with single A mismatch being the highest, with a mean percentage of 12.4% (**Fig 6c**, right).

**Fig 6:**
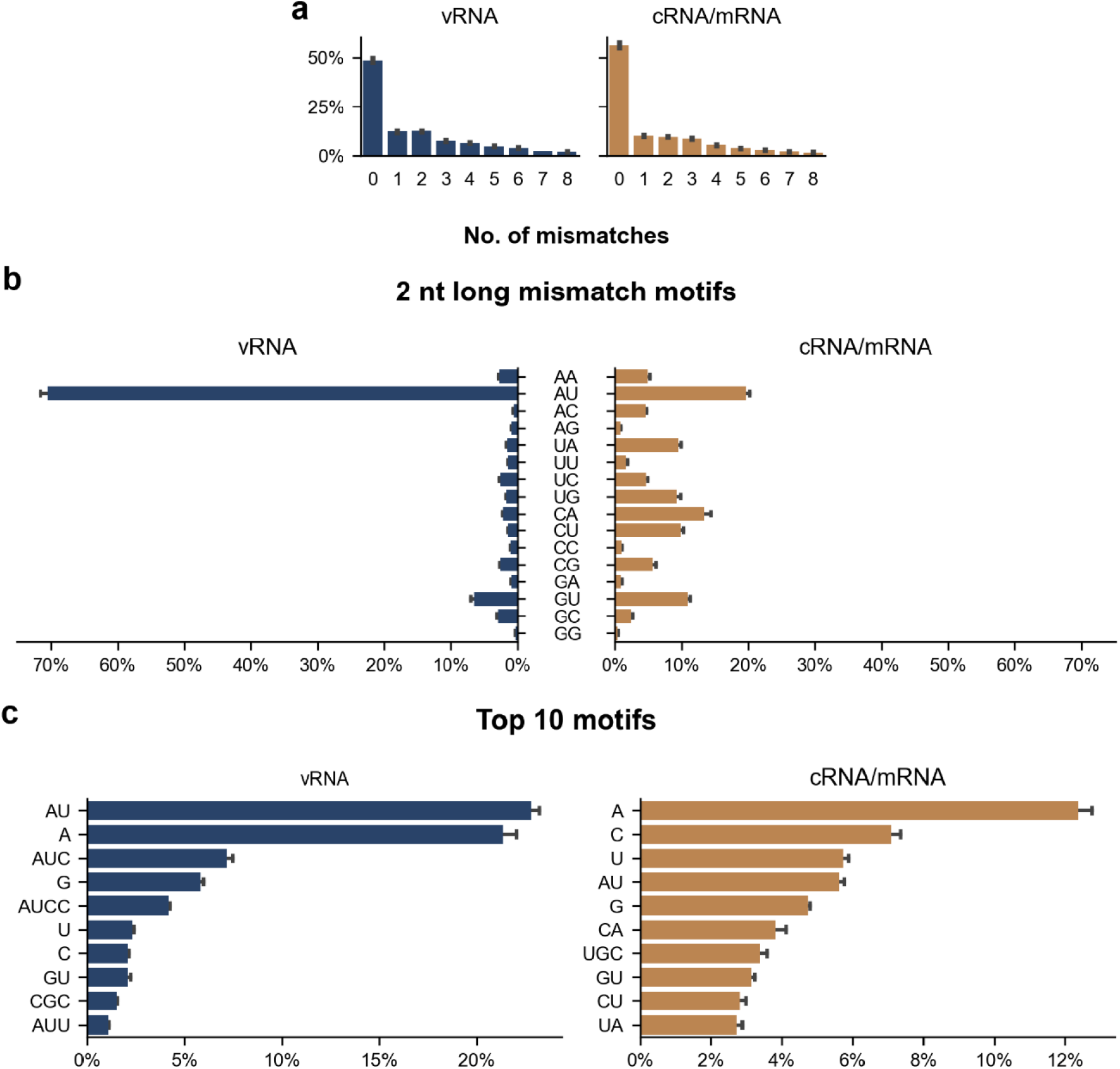
High Misincorporation Rates Associated with Pausing at the 3’end of Nascent RNA transcripts. Incorporation of mismatched nucleotides at the 3’ end during IAV RNA synthesis (n = 3, mean ± SEM). **a.** Percentage of mismatched nucleotides at the 3’ end of vRNA (left) and cRNA/mRNA (right). A value of 0 indicates reads with no mismatches at the 3’ end. **b**. Percentage of 2-nucleotide long motifs at the 3’ end of vRNA (left) and cRNA/mRNA (right). **c**. The top 10 mismatch motifs at the 3’ end of vRNA (left) and cRNA/mRNA (right), ranked by mean percentage.

## Discussion

Viruses depend on the RdRp complex to replicate and transcribe their genomes. While structural and enzymology studies have characterized RdRp activity in isolated environments, a direct connection to physiological setting is missing. To address this, it is important to study the genome-wide dynamics of RdRp during the replication and transcription of the entire viral genome within infected cells using nascent RNA sequencing. However, there is an absence of technology to access and analyze viral nascent RNA species that habour crucial mechanistic information. By investigating the dynamics of replication and transcription in host cellular environment through analyzing this undiscovered nascent RNA species, we can unravel the molecular mechanisms underlying viral RNA synthesis and identify new targets for developing antivirals.

We introduce Total Elongating Nascent VIral Polymerase single-molecule Sequencing (TenVIP-seq), the first platform designed to study nascent viral RNA synthesis at single-nucleotide resolution across entire viral genomes. By leveraging this comprehensive and unbiased (PCR-free) approach, we discovered and analyzed nascent viral RNA species and uncovered transient RdRp behaviors that are critical to understanding key regulatory mechanisms. These behaviors include polymerase pausing, termination signals, and the RdRp’s responses to mutations, as well as the effects of antivirals and host factors on RdRp activity in a genuine genome-wide setting. Our findings lay the foundation for future research into viral genome regulation, the impacts of antiviral impacts, adaptation, evolution, and virus-host interactions, all with high-resolution, genome-wide insights.

Existing nascent RNA sequencing methods for eukaryotes and prokaryotes are not adaptable to study viral RNA synthesis. For instance, viral RdRp does not efficiently incorporate biotin-NTPs^64^, which are essential for Pro-Seq. Furthermore, IAV PB2 antibodies do not efficiently capture the viral genomic (vRNA) and anti-genomic (cRNA/mRNA) species needed for analysis. Ensuring that purified viral RNA sequences are nascent and exclusively produced by the viral RdRp present additional challenges^64^. To address these limitations, we engineered a customized nascent RNA sequencing technology platform specifically designed for viruses. We generated an IAV strain expressing a Strep-tagged PB2 subunit, enabling selective enrichment of actively transcribing viral polymerase complexes from the nuclei of infected host cells. Critically, we introduced a novel adaptor ligation approach that enhances the efficiency of adaptor attachment to transient nascent RNA species and enables direct sequencing of not only mRNA, but also cRNA and vRNA. By employing single-molecule sequencing, we achieved accurate identification of the last nucleotide at single base-pair resolution and detect rare, transient misincorporation events. The authenticity of the nascent viral RNA sequences was confirmed through multiple lines of evidence.

We present the first pausing landscape of a virus, highlighting the significance of pausing in various biological phenomena. This work allows us to discover potential pausing sites of the viral RdRp during both replication and transcription, a previously elusive aspect of virus biology. The first question we seek to address is to determine whether viral polymerase pausing is a genuine biological phenomenon observable in cellular environments, with reproducible pausing sites at specific genomic loci. Importantly, these pausing signatures were absent from a control dataset of mature viral RNAs, confirming that the observed patterns reflect the true dynamics of the viral replication and transcription processes within the host cell. A longstanding question in virology is how viruses maintain a delicate balance between genome replication and gene expression^65^, all utilizing a single multifunctional enzyme, the viral RdRp. Analysis of the 3’ end positions of the nascent RNA reads using TenVIP-seq uncovered distinct pausing profiles for vRNA and cRNA)/mRNA strands, providing new insights into the regulatory mechanisms underpinning viral RNA synthesis.

To investigate whether specific genomic sites exist where the viral polymerase exhibits a higher probability of pausing suggestive of a putative regulatory site, we explored the role of RdRp pausing in regulating genome replication and transcription. Our findings indicate that viral RdRp pausing plays a critical role in regulating mRNA transcription. Using TenVIP-seq, we identified peaks of RdRp pausing at the poly(U) track polyadenylation sites of each viral segment during mRNA transcription, with the most pronounced pausing occurring at the second U in the 5-6 nucleotide poly(U) track. This behavior suggests that when the RdRp encouteres the poly(U) tract on the vRNA template, it pauses while potentially incorporating the corresponding adenine residues to synthesize the poly(A)-tail of the mature viral mRNA.

TenVIP-seq provides a novel approach to understanding how antiviral drugs like NTP analogs influence RdRp activity across viral genomes. NTP analogs, such as T-705, have been utilized in the treatment of influenza viruses and COVID-19, yet their effects on viral RdRp transcription across complete viral genomes have largely remained unexplored. Our study offers new insights into the mechanisms of action of antiviral nucleotide analogs through TenVIP-seq. Interestingly, TenVIP-seq revealed that T-705 predominantly increased the intensities of existing pausing sites on the viral genome rather than introducing new pausing sites. By selectively enhancing the frequency or duration of pausing at specific genomic locations, T-705 may disrupt the normal kinetics and processivity of the viral polymerase. This work represents the first effort to elucidate the mechanisms of an antiviral drug targeting RdRp directly at the genomic level. This is significant as it suggests that the drug targets core aspects of viral replication and transcription, rather than inducing drastic effects such as complete polymerase inhibition or creating insurmountable pausing sites. Modulating existing pausing behavior may be a more nuanced yet effective strategy to impair viral genome synthesis and proliferation. This systems-level understanding of the drug’s impact on the viral polymerase complex and its dynamics across the entire viral genome offers valuable insights that can inform the development of more effective antiviral strategies. Future work may involve time-dependent pausing profiling, offering a more physiologically relevant platform than traditional enzymology assays to rapidly screen nucleotide analogs and predict their efficacy against viral RdRp.

Host factors play a critical role in virus replication, yet their effects on RdRp transcription and replication have not been measured at genomic resolution within cells. This limitation has hindered progress in identifying and understanding how host factors influence viral polymerase activity in a cellular and genomic environment. Current methodologies often failed to capture the *in vivo* impacts of host factors on RdRp activity, as *in vitro* data do not always translate to the physiological context, and endpoint virus titer assays provide only limited insights. Our study demonstrates the potential of TenVIP-seq to offer a systems-level perspective, elucidating how cellular proteins or other host factors alter RdRp processivity, pausing, or termination events during viral replication and transcription. As a case study, we investigated the role of the host factor TRIM25, an E3 ubiquitin ligase essential for the innate immune sensing of influenza virus and the induction of type I interferons. Previous studies have indicated that TRIM25 restricts influenza virus replication. However, the underlying mechanisms remain incompletely understood. Using TenVIP-seq, we compared the pausing profiles of the influenza RdRp in wild-type and TRIM25-knockout A549 cells. Our analyses revealed that the knockout of TRIM25 led to a remodeling of the viral genomic pausing landscape, with a global reduction in the intensity of pausing sites across the influenza genome. This suggests that TRIM25 plays a critical role in shaping the replication and transcriptional dynamics of the viral RdRp across the entire viral genome. Specifically, TRIM25 appears to slow viral replication by increasing the frequency and intensity of RdRp pausing on a genome-wide scale. This study provides the first genomic-resolution observation of a host factor recalibrating viral RdRp function. The observed alterations in RdRp pausing patterns in the absence of TRIM25 likely have important functional implications for the coordination and regulation of viral genome replication and gene expression. These results highlight the value of our TenVIP-seq approach in providing high-resolution insights into how host factors modulate the dynamic behavior of viral replication machinery at the genome level. Insights gained from TenVIP-seq studies of factors like TRIM25 could uncover new drug targets at the virus-host interface.

Nascent viral RNA sequencing provides valuable insights into the mechanisms underlying polymerase fidelity during viral transcription and replication. While it is well established that DNA polymerases and eukaryotic RNA polymerases with proofreading capabilities can backtrack and pause following misincorporation events, the behavior of viral polymerases lacking such proofreading functions remains less understood. To address this knowledge gap, we employed the high-resolution TenVIP-seq approach to investigate whole-genome polymerase dynamics during viral RNA synthesis. Strikingly, our analyses revealed that 51.6% of vRNA and 44.0% of cRNA/mRNA reads contained misincorporated nucleotides at the 3’ end, indicating that polymerase errors can lead to stalling even in the absence of proofreading. The presence of misincorporations at the 3’ terminal near pausing sites suggests a higher mutational rate than previously reported, which may have important implications for the evolution of influenza viruses. Misincorporations during viral RNA synthesis can introduce genetic diversity and potentially facilitate the emergence of new viral variants or quasispecies.

We also compared the genomic motifs where IAV RdRp pauses and found that RdRp exhibits differential pausing preferences for replication versus transcription. Further investigation with RdRp mutants, such as the β-hairpin of PB1 with proline 651, which is critical for efficient RNA synthesis during replication but not necessary for internal initiation and transcription^66^, could provide insights into how RdRp navigates between replication and transcription. The distinct motif preferences for pausing during vRNA versus cRNA and mRNA synthesis underscores the complex interplay between polymerase dynamics, template sequence, and transcriptional regulation.

While TenVIP-seq has significant advantages, there are areas for improvement. The inability to identify reads shorter than 15 nt limited our analysis of the vRNA and cRNA promoters, which consist of 9-12 nt at the 5’ end. Also, the current preparation does not separate cRNA from mRNA. Future optimization is needed to address these issues. Nevertheless, the TenVIP-seq approach holds transformative potential for advancing our understanding of host-virus interactions and accelerating antiviral discovery. As an unbiased, genome-wide technique that closely resembles the physiological environment compared to traditional enzymology assays, TenVIP-seq can provide critical mechanistic insights that were previously inaccessible. We demonstrated its utility in identifying regulatory sites affecting RdRp dynamics in RNA synthesis and have made resource and data publicly available for researchers to study novel regulatory elements. By elucidating how host factors like TRIM25 modulate the transcriptional dynamics of viral RdRp, including the regulatory sites for the viral polymerase element, this method can reveal the molecular determinants governing zoonotic transmission and host adaptation of emerging viruses. Our findings also indicate that genome replication and transcription exhibited distinct pausing and misincorporation profiles by the same RdRp, paving the way for future research into how viral and host factors may regulate differently these two processes.

Nascent RNA sequencing has opened transcription dynamics *in vivo* as a research field and driving novel biological insights, as it is becoming a fundamental tool for studying gene regulation. For instance, nascent RNA sequencing provides valuable insights into how eukaryotic DNA-dependent RNA polymerases such as RNA polymerase II coordinate and regulate gene expression at the genomic level. By tracking the synthesis of nascent viral RNAs, specific mechanisms and regulatory strategies employed by the RNA polymerase II and other host factors to control these fundamental viral processes of transcription can be elucidated^18,43–54^. This allows for unparalleled temporal observations of the dynamic nature of the transcriptome and the regulatory mechanisms at work within living cells^54,55^. By selectively capturing newly synthesized and unprocessed RNA molecules, theses powerful approaches provide a precise and comprehensive understanding of gene regulation and expression, surpassing the limitations of traditional RNA sequencing^56^. Existing nascent RNA sequencing techniques were developed for eukaryotic and prokaryotic systems^43–47,52,53^. For example, native elongating transcript sequencing (NET-seq) uses immunoprecipitation on the RNA polymerase II to capture nascent transcript, which are the newly synthesized RNA molecules still attached to the elongating RNA polymerase complex^43,45,57^. Precision Nuclear Run-On sequencing (PRO-seq) utilizes biotin-labeled nucleotide analogs such that transcription is halted to provide nucleotide-resolution mapping of nascent RNA transcripts^52,58^. Recently, nano-COP utilizes Nanopore direct RNA sequencing (DRS) to observe splicing events in nascent transripts^59,60^. DRS as a PCR-free method offers the advantage of assessing splice variants as single RNA molecule and unbiased ratio^35^. In eukaryotic systems, the application of nascent RNA sequencing has uncovered the dynamic transcription of enhancer regions, leading to the identification of enhancer RNAs and their pivotal roles in gene regulation^46,48^. Moreover, this technique has shed light on the intricate process of co-transcriptional splicing, as well as the regulatory functions of transcriptional pausing^50,61,62^. In bacteria, nascent RNA sequencing has proven instrumental in the discovery of novel regulatory elements, such as small RNAs and riboswitches, and has facilitated the comprehensive characterization of transcriptional responses to environmental changes and stress conditions^49,51,63^. This enhanced understanding of bacterial transcriptional dynamics holds immense potential for advancing our knowledge of microbial physiology and adaptability, as well as opening new windows of opportunities for the development of next generation antibiotics. Learning from nascent RNA sequencing in eukaryotes, questions remain about the regulation of viral genome replication and transcription offers an opportunity to finally be addressed, with transformative insights, such as mechanisms of viral gene replication, understanding polymerase pausing and dynamics, antiviral drug inhibition and resistance mechanisms, host-virus interactions viral polymerase mutation, impact of sequence context, temporal dynamics of RNA synthesis, study of critical regulatory elements, differential impact on subgenomic transcripts, comprehensive view of viral life cycle, among others.

## Methods

### Cell lines and chemicals

Human embryonic kidney cells 293T (HEK293T; ATCC), human alveolar basal epithelial cells A549 (Fenghui Biotechnology) and A549 TRIM25 Knockout (A549 TRIM25-KO; Ubigene) cells were cultured in Dulbecco’s modified Eagle’s medium (DMEM; Invitrogen), supplemented with 10% fetal bovine serum (FBS; Invitrogen) and 1% penicillin-streptomycin (P/S) (Invitrogen). Madin-Darby canine kidney (MDCK; Fenghui Biotechnology) cells were cultured in Advanced DMEM (Invitrogen) supplemented with 4% FBS, 4 mM L-alanyl-L-glutamine (Macklin) and 0.4% P/S (Invitrogen). Cells were cultured at 37 °C, 5% CO_2_ in a humidity-controlled incubator.

Favipiravir (T-705; Macklin) was prepared at a stock concentration of 15 mM in RNase-free water.

### Virus generation, propagation, and titration

The wild-type influenza A/WSN/1933(H1N1) and recombinant PB2-strep viruses were rescued by reverse genetics. The plasmids pHW2000-PB2/PB1/PA/NP/HA/NA/NS/M described by Hoffmann et al.^67^ were used for virus rescue. The plasmid pHW-PB2-strep was constructed by orderly fusing the twin-strep tag coding sequence and the 5’ terminal 143 nucleotides of the PB2 segment to the last amino acid of the PB2 open reading frame of pHW-PB2 plasmid using NEBuilder HiFi DNA Assembly Cloning Kit (New England Biolabs) as previously described^68^. MDCK and HEK293T cells were co-transfected by the 8 pHW2000-WSN plasmids in a 6-well plate. Cells at 80% confluency in each well cultured in the virus infection medium (DMEM supplemented with 20 mM HEPES (Gibco) and 1 µg/ml TPCK-trypsin (Sangon Biotechnology)) were transfected with 250 ng of each pHW2000 plasmid with 8 µg Polyethylenimine “Max” (PEI-MAX; Polysciences) in 200 µl Opti-MEM medium (Gibco). The culture media were collected and spun down after 72 hours. The supernatants were flash-frozen in liquid nitrogen and stored at -80 °C. The recombinant virus was propagated in A549 cells at the M.O.I. of 0.005 in the infection medium.

Viral titer was measured by standard plaque assay in MDCK cells. In brief, a 12-well plate with 100% confluency MDCK cells was washed twice with 1× PBS. Cells in each well were then inoculated with 300 µl of the recombinant virus stock at 10-fold serial dilution in the infection medium. The virus inoculum in each well was removed after 1 hour of incubation at 37 °C. The infected cells were covered by 1 ml of the infection medium supplemented with 0.8% low melting-point agarose (Invitrogen) and incubated at 37 °C with 5% CO_2_. After 3 days of incubation, the agar medium was taken off, and the infected MDCK cells were fixed with 70% ethanol for 10 minutes and stained with 2% crystal violet solution (BBI). The crystal violet solution was discarded before cell washing with nuclease-free water, and the plaque was counted to calculate the titer.

For viral growth curves, a 12-well plate of confluent A549 cells were infected by the virus at an M.O.I. of 0.005 in triplicates for each virus. The supernatant of the sample was collected 3, 7, 10, 24, and 36 hours after infection, and stored in -80 °C before the TCID50 assay. The TCID50 assays were done by first preparing a 96-well plate with 100% confluency MDCK cells that were washed twice with 1× PBS. The supernatant of each timepoint was then serial diluted 5-fold from 10^-2^ in the virus infection medium. Cells were covered by 100 μl of the diluted supernatant and incubated at 37 °C with 5% CO_2_. After 3 days of incubation, the cells were washed with PBS and fixed with 70% ethanol and stained with 2% crystal violet. The titer of each sample was calculated using the Spearman–Kärber Method^69^.

### Data preprocessing and analysis

Outputs from sequencing as fast5 files were first converted to pod5 files for basecalling. The trimmed fastq files were then aligned to a custom reference consists of the human transcriptome the assembly GRCh38.p14 from Ensembl, the A/WSN/1933(H1N1) viral genome^70^, the yeast ENO2 transcript from Ensembl which constitutes RCS, and the ERCC spike-in reference using minimap2^71^. The unmapped reads from the output were then aligned again for short reads. The output sam files were concatenated together for downstream analysis. Reads with poly(A)-tail, quality < 1, and not uniquely mapped were identified and filtered. The remaining mapped reads were categorized as cRNA/mRNA or vRNA based on the read direction. The coverage, sequencing depth, the last nucleotide position, and statistical analysis were performed using samtools 1.20^72^ and Python 3.12.4^73^.

### Western blot analysis

To assess the purity of subcellular fractionation, equal amounts of total protein from cytoplasmic and nucleoplasm fraction were incubated in Laemmli buffer at 95 °C for 5 min, and then subjected to gel electrophoresis. The proteins separated on SDS-PAGE (15% for Histone H3, 8% for the others) were then transferred to a PVDF Membrane (Merck) by using Power Blotter XL System (Invitrogen), which was blocked with Protein-Free Blocking Buffer (P0242; Beyotime). The membrane was incubated with a primary Histone H3 Rabbit pAb (1:8000 dilution; A2352; ABclonal), β-Tubulin Rabbit pAb (1:5000 dilution; AC008; ABclonal), Strep II-Tag Mouse mAb (1:20000 dilution; AE066; ABclonal) and Influenza A PB2 pAb (1:5000 dilution; PA5-32220; Invitrogen) overnight at 4 °C. The membrane was washed three times with TBST (0.1%), then incubated with Goat anti-Rabbit IgG Secondary Antibody, HRP (1:40000 dilution; A16110; Invitrogen), or Donkey anti-Mouse IgG Secondary Antibody, HRP (1:20000 dilution; A16017; Invitrogen) for 1 h at room temperature. The blot was detected by chemiluminescence using SuperSignal West Atto Substrate (A38554, Thermo Scientific) at room temperature, and then imaged with Odyssey M imaging system (LI-COR).

Primary TRIM25 Rabbit pAb (1:2000 dilution; 12573-1-AP; Proteintech) was used when testing the expression of TRIM25 protein in A549 WT and TRIM25-KO cells. 1 × 10^6^ cells were lysed in 150 mM NaCl, 50 mM Tris pH 7.4, 1% NP-40, 2.5 mM MgCl_2_, 5 mM EDTA, 1 mM DTT, 1× PIC. 20 μg protein from lysed cell supernatant was resolved by using an 8% SDS-PAGE gel and the blot analysis proceeded as described above.

### RT-qPCR

To compare the polymerase activity between the wild-type and Strep-PB2 virus, 1 × 10^6^ A549 cells were infected by two viruses respectively at the M.O.I of 1. The total RNA was extracted 4 hours post-infection using NucleoZOL as described above. 500 ng RNA was reverse transcribed by using HiScript III 1^st^ Strand cDNA Synthesis Kit (R312-02; Vazyme) with a 0.4 μM primer mix specific^74^ for both NP vRNA and cRNA, and oligo-dT (R312-02; Vazyme) for host mRNA. The reaction mixture was incubated at 50 °C for 15 min and terminated at 85 °C for 5 s. 1 μl of cDNA was added to the PCR reaction mixture by using Taq Pro Universal SYBR qPCR Master Mix (Q712-00; Vazyme) with 0.4 μM forward and reverse primer. The PCR reaction was performed on a real-time PCR detection system (CFX96; Bio-Rad) under follow conditions: 95 °C for 30 s, followed by 45 cycles of 95 °C for 10 s and 60 °C for 30 s. Data were normalized to ACTB gene expression and fold changes in gene expression relative to 0 hours post-infection were calculated using the 2^−ΔΔCT^ method with all experiments repeated in triplicate.

To investigate the impact of the T-705 on newly synthesized RNA expression, different concentrations of the drug were added to 1 × 10^6^ A549 cells 5 hours before infection and maintained for the following 4 h infection at M.O.I. of 1. The level of newly synthesized vRNA and cRNA of NP were measured as described above and normalized to GAPDH expression.

## Data availability

All sequencing data are available at NCBI BioProject ID PRJNA1161490.

## Acknowledgment

We are grateful to the current and former members of the Cheung Laboratory for discussions and critical review of the manuscript. We thank the Li Ka Shing Translational Omics Platform for technical support. This work was supported by Collaborative Research Fund (C4032-21G) and Early Career Scheme (24303324) for P. P. C., National Natural Science Foundation of China/Research Grants Council of Hong Kong Joint Research Scheme (N_CUHK484/23) for P. P. C. and X. X. The Hong Kong RGC Postdoctoral Fellowship Scheme (PDFS2324-4S06) for X. X.

