## supplemental file for "Genome-wide Viral Nascent RNA Sequencing Unveils Polymerase Pausing Landscape at Single-nucleotide Precision"

### Supplemental figures and data

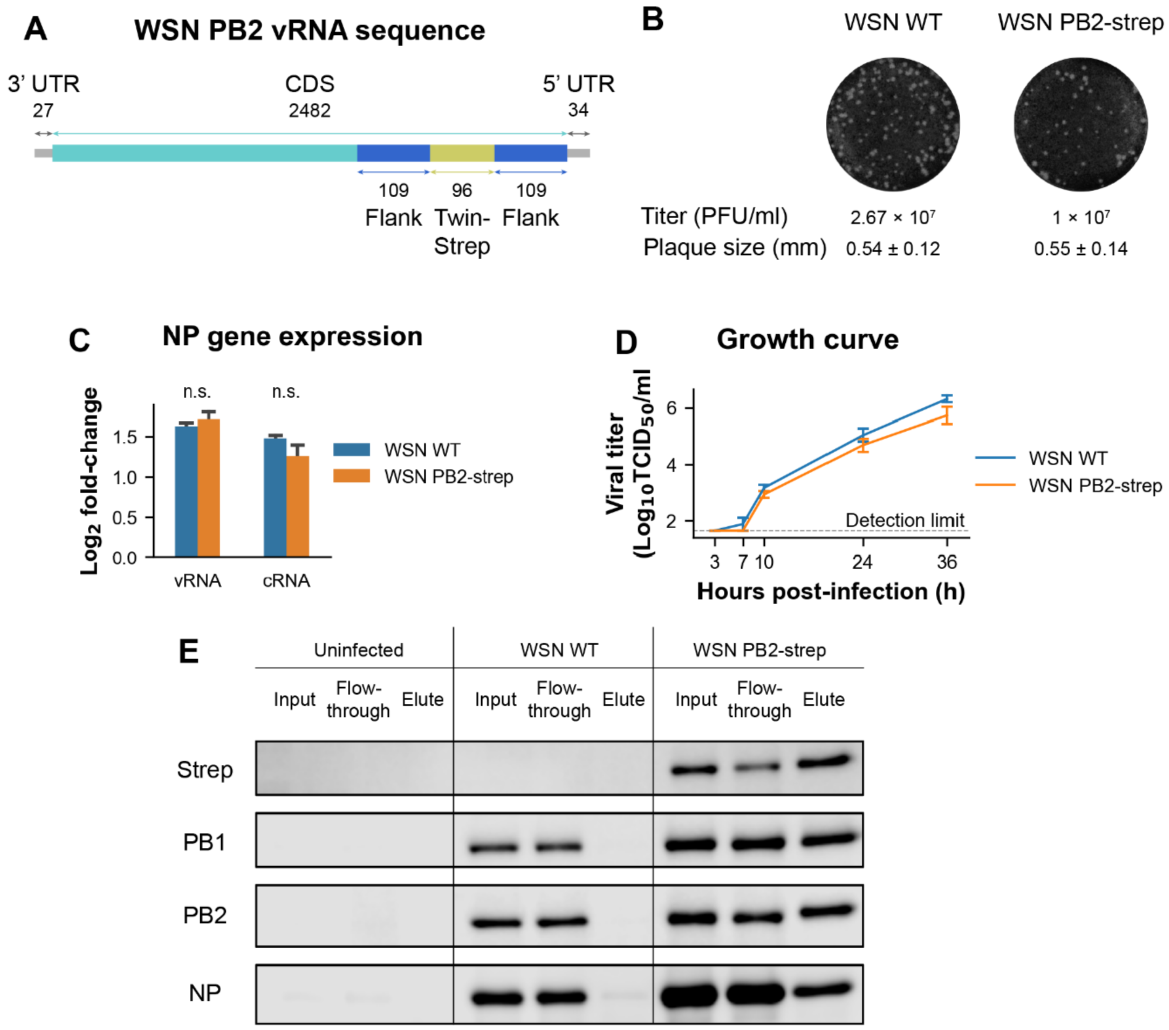


**Fig. S1**: Construction of Recombinant Influenza A Virus with a Functional Strep-Tag at the PB2 Subunit. **a.** Sequence of the PB2 vRNA construct. **b.** Plaque assay results comparing wild-type WSN and PB2-recombinant WSN. **c.** Real-time quantitative PCR results comparing NP expression levels between wild-type WSN and PB2-recombinant WSN. **d.** Viral growth curve derived from viral titration measured by $TCID_{50}$. **e.** Western blot analysis showing antibodies against Strep-tag, PB1, PB2, and NP in the input, flow-through from Strep bead binding, and final elution prepared for downstream library construction.

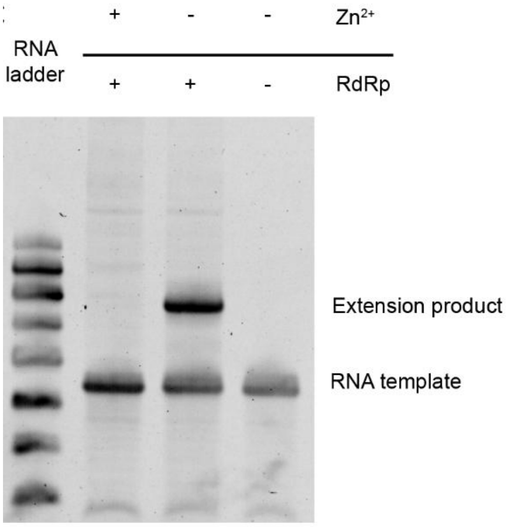


**Fig. S2**: Urea PAGE gel image of the in vitro transcription (IVT) assay using recombinant IAV polymerase in the presence of $Zn^{2+}$ at 30 °C for 1 hour.

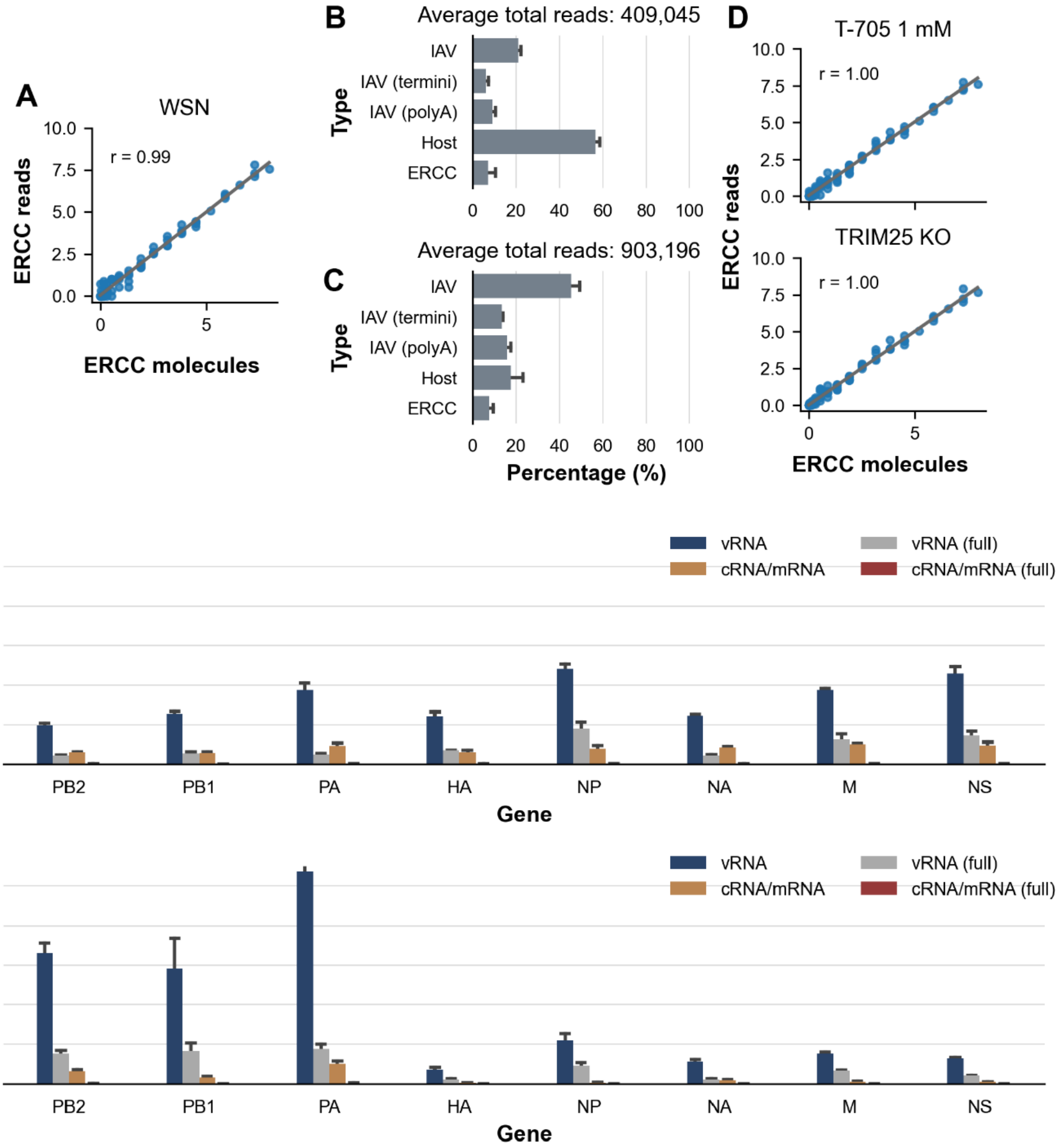


**Fig. S3**: Read Depth and Dynamic Range of TenVIP-seq. **a.** Assessment of the dynamic range of sequencing data from WSN-infected A549 cells using the ERCC spike-in mix (Invitrogen). Pearson's correlation coefficient = 0.99. **b.** Percentage of each RNA species in uniquely mapped reads from A549 cells treated with T-705 prior to WSN infection. Data presented as mean ± SEM, n = 3. **c.** Percentage of uniquely mapped reads in TRIM25 knockout (KO) A549 cells. Data presented as mean ± SEM, n = 3.c. **d.** Dynamic range assessment of sequencing data from T-705-treated and TRIM25-KO cells. **e, f.** Percentage of vRNA and cRNA/mRNA reads for each IAV gene, including reads aligned to the 3' end of the respective RNA species: **e.** T-705 treatment, n = 3. **f.** TRIM25-KO cells, n = 3.

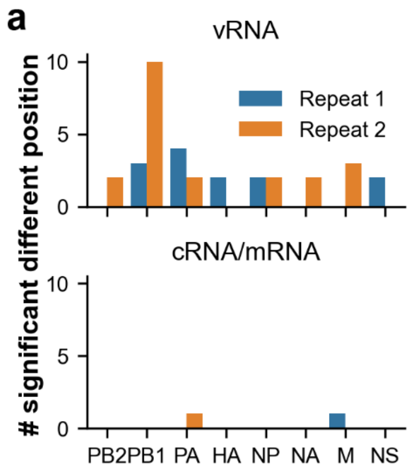


**Fig. S4:** Biological Replicates of TenVIP-seq Show Minimal Differences. **a**. Number of significantly different positions identified by t-test. Top: vRNA; Bottom: cRNA/mRNA.

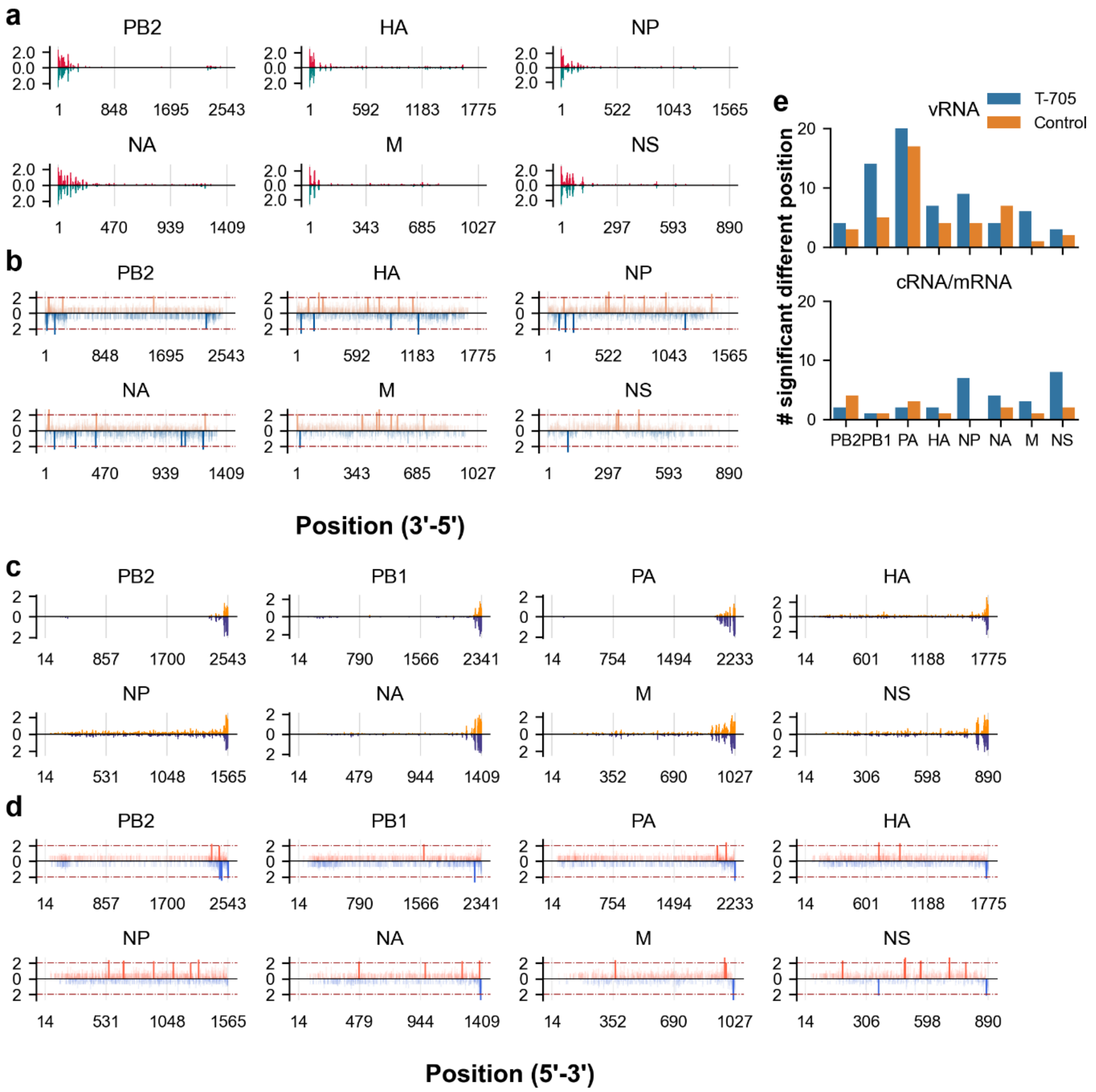


**Fig. S5.:** T-705 Increases RdRp Pausing Frequency. **a**. Comparison of RdRp positions during vRNA synthesis of PB2, HA, NP, NA, M, and NS between T-705-treated A549 cells and control cells infected with WSN at a multiplicity of infection (M.O.I.) of 1 (n = 3). **b**. Differences in vRNA single nucleotide coverage between T-705-treated and control cells, visualized by -$\log_{10}$($p$-value) following Student's t-test. Dashed line indicates a $p$-value of 0.01. **c**. Comparison of RdRp positions during cRNA/mRNA synthesis for all WSN genes between T-705-treated and control cells. **d**. Differences in cRNA/mRNA single nucleotide coverage between T-705-treated and control cells. **e**. Number of significant differences identified by t-test. Top: vRNA; Bottom: cRNA/mRNA.

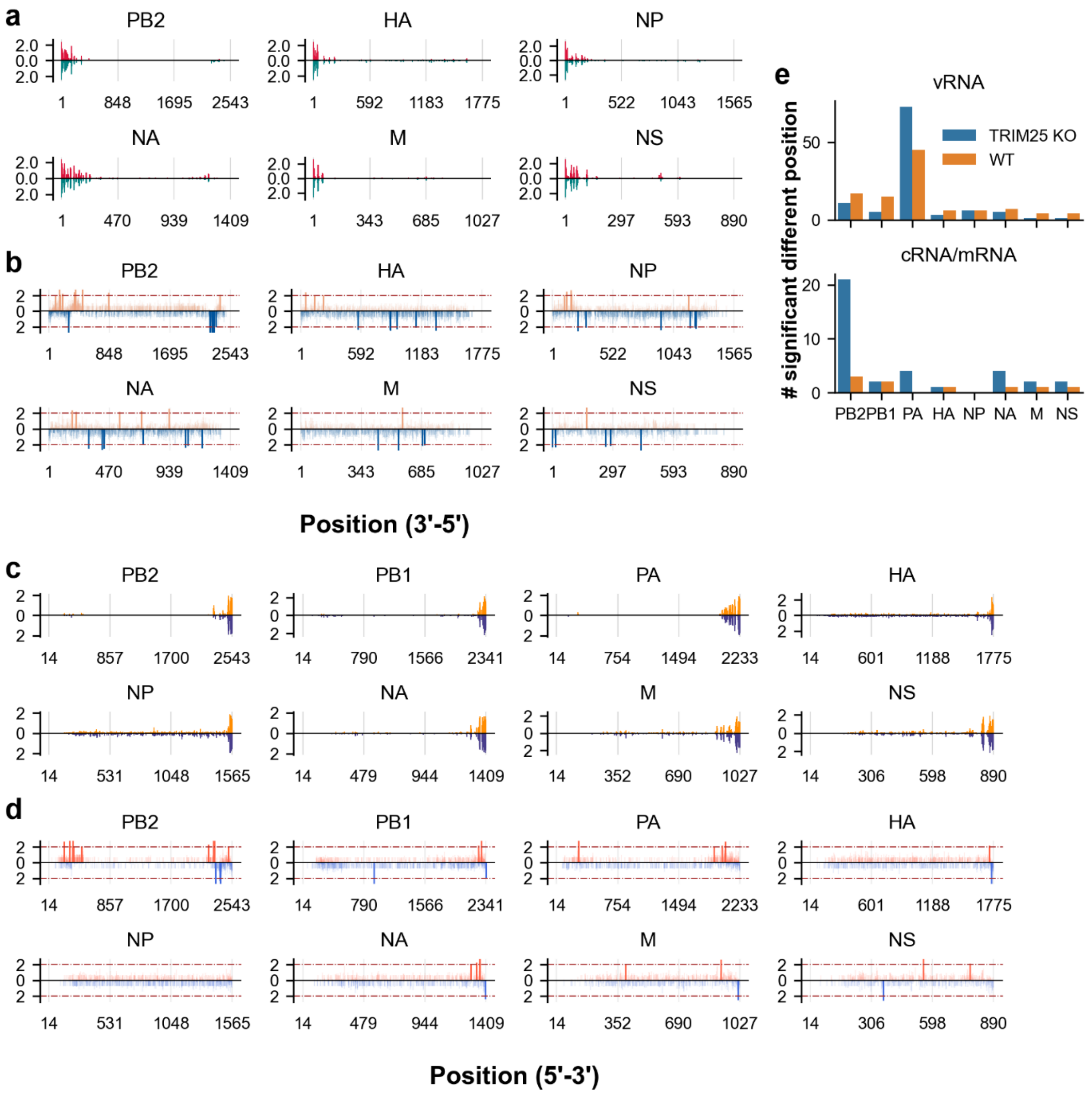


**Fig. S6:** TRIM25-KO alters pausing profile. **a.** RdRp position comparison during vRNA synthesis of PB2, HA, NP, NA, M and NS between TRIM25-KO and wild- infected by WSN at M.O.I. of 1, n = 3. **b.** Differences of the vRNA single nucleotide coverage between TRIM25-KO and WT, visualized by the -log(*p*-value) after Student's t-test. Dashed line indicates *p*-value at 0.01. **c.** RdRp position comparison during cRNA/mRNA synthesis of all WSN genes between TRIM25-KO and WT. **d.** Differences of the cRNA/mRNA single nucleotide coverage between TRIM25-KO and WT. **e**. Number of significant differences from t-test. Top: vRNA; bottom: cRNA/mRNA.

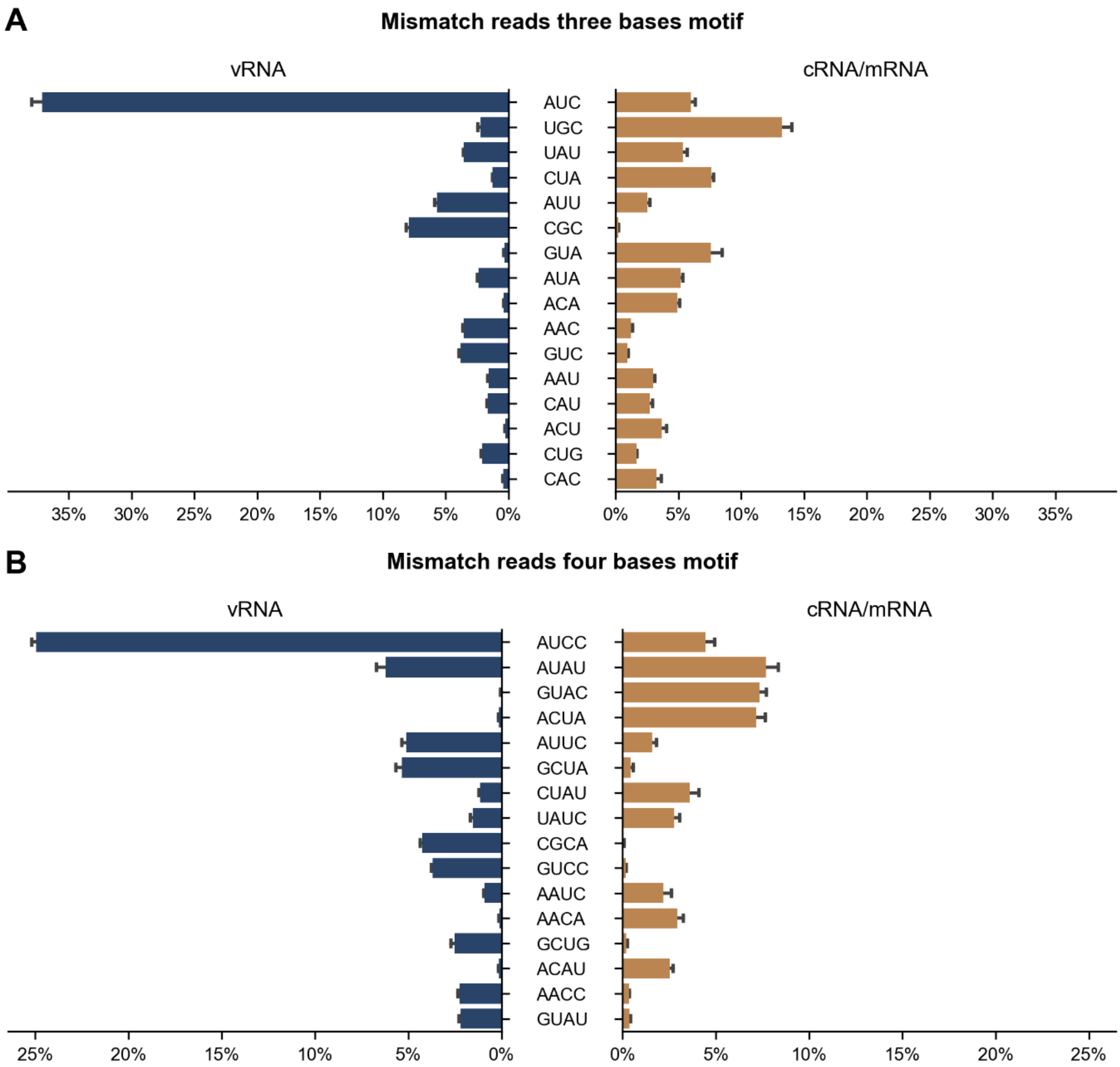


**Fig. S7:** 3' End Mismatch Motifs. 10 motifs with the highest mean percentage were shown. **a**. Percentage of mismatched 3-nucleotide long motifs at the 3' end of vRNA and cRNA/mRNA. **b**. Percentage of mismatched 4-nucleotide long motifs at the 3' end. Data are presented as mean ± SEM, n = 3.
